## Supplementary Figures for "Integration of single cell gene expression data in Bayesian association analysis of rare variants"

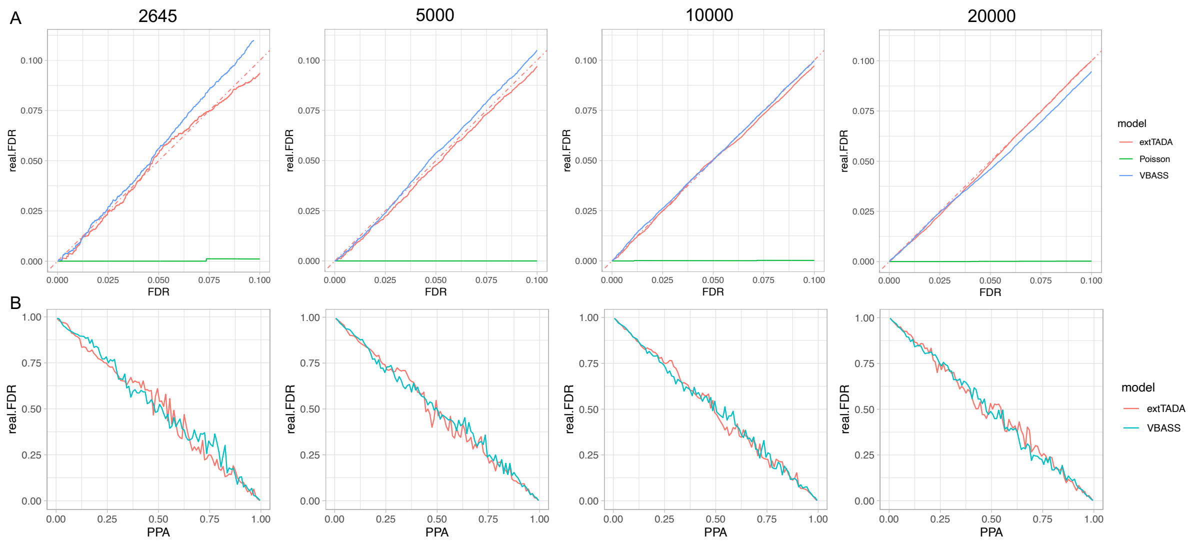


Supplementary Fig. 1. A) False discovery control for two models and Poisson test, only show the part with FDR ≤ 0.1. X-axis, estimated false discovery rate (FDR) from the model, y-axis, real false discovery rate in simulation. B) Local false discovery control for two models. X-axis, posterior probability attribute (PPA) from the two model, y-axis, real false discovery rate in simulation. Each dot represents a gene set with 100 genes with close PPA.
